# Flexible self-protection as evidence of pain-like states in house crickets

**DOI:** 10.1101/2025.09.12.675781

**Authors:** Oscar Manzi, Kate E. Lynch, Daniel M. Allman, Tanya Latty, Thomas E. White

## Abstract

The possibility that insects experience pain is a frontier question at the intersection of behaviour, cognition, and philosophy of mind. Interest has been fuelled not only by anatomical discoveries but also by expanding behavioural and comparative evidence. Leading frameworks emphasise behavioural indicators of pain-like experience such as flexible, targeted responses to harm beyond reflexive withdrawal. Here we tested for such responses in the house cricket (*Acheta domesticus*), a species of evolutionary and commercial importance. Using a fully blinded, within-subjects design, we applied noxious heat, innocuous tactile contact, or no contact to a single antenna under lower- and higher-stress environmental conditions and recorded grooming behaviour. Crickets were significantly more likely to groom the noxiously stimulated antenna, and did so for longer, than under control or tactile treatments. Grooming also showed a distinct temporal profile, with elevated activity sustained across the early observation period. Environmental condition and sex had no effect, indicating that self-protective grooming was expressed consistently throughout. These findings provide robust evidence of flexible, site-directed self-protection in Orthoptera, addressing a key gap in evidence for pain-like states outside vertebrates. This strengthens the case for consideration of insect welfare and bears on how felt experience is distributed across the animal kingdom.

## Introduction

Animals navigate a world of shifting threats and fleeting opportunities, where thriving depends not just on reacting, but on evaluating. Sentience is commonly defined as the capacity for subjective, valenced experience, and is increasingly (but not universally; Eisemann et al., 1984) argued to be a core, widely-distributed biological capacity that shapes how animals respond to the world (Low et al., 2012). Under adaptive accounts, it is variously hypothesised to support deep forms of learning (Ginsburg & Jablonka, 2019), inform complex decision-making (Veit, 2023), and enable flexible, goal-directed behaviour (Elwood, 2011). While its neural substrates likely differ widely across taxa, a c role—enabling animals to evaluate and respond to salient internal and external conditions—appears broadly conserved (Birch et al., 2021; Gibbons, Crump, et al., 2022; Veit, 2023). In both humans and non-human animals, felt experiences are suggested to enable responses that go beyond ‘mere reflex’ or stereotyped output into behaviours marked by complexity, context sensitivity, and/or specificity (Ginsburg & Jablonka, 2019; Veit, 2023). Yet, because experience is inherently private, its presence is difficult to infer, especially in invertebrates where brains lack close homology to mammals and behavioural profiles can appear equally unfamiliar, resisting easy comparison to vertebrate models.

Pain presents a critical test case. Distinct from nociception—the unconscious detection of damaging stimuli—pain is typically (though not uniformly) defined as a negatively valenced experience of harm (Raja et al., 2020; Sneddon et al., 2014). Unlike nociceptive reflexes, which can occur without complex central processing, pain supports persistent behavioural change, lasting avoidance, and context-sensitive self-protection (Elwood, 2019; Sneddon et al., 2014). Nociception is widespread across the animal kingdom and confers clear evolutionary value, but pain is thought to involve additional costs such as the metabolic demands of centralised processing and the risk of behavioural distraction (P. Bateson, 1991). Such costs imply that pain would not evolve without compensating benefits. From this perspective, pain can be seen as an adaptation that builds upon nociception to enable richer forms of behaviour and learning and, ultimately, support long-term fitness.

Efforts to assess pain and sentience in animals have changed markedly over time. Historically, the capacity for subjective experience was tightly coupled to anatomical similarity with humans, and the scope of morally considerable animals was narrowly drawn. That boundary has since shifted outward, with birds, fish, and more recently cephalopod molluscs and decapod crustaceans recognised as potentially sentient (Birch et al., 2021; Gibbons, Crump, et al., 2022; Sneddon, 2019). This creeping frontier reflects a broader shift in thinking: rather than asking which brains are most like ours, a question with clear failings (Panksepp, 2016), researchers now ask which behavioural and physiological features justify the inference of experience (Crump et al., 2022; Jablonka & Ginsburg, 2022; Sneddon et al., 2014). Frameworks such as that proposed by Crump et al. (2022) have codified this shift by outlining eight empirically testable criteria for pain-like states and, by inference, sentience, including motivational trade-offs, associative learning, and flexible self-protection. While the framework, understandably, does not provide a formal method for balancing these criteria — though subsequent efforts have begun to address this (Fischer et al., 2025; Gibbons & Chittka, 2022) — it highlights how different lines of evidence can be assembled to guide judgement about the likelihood of pain experience.

In this context insects have emerged as a critical frontier. Once dismissed as too small-brained or simple to support experience (Eisemann et al., 1984), they are now known as capable of remarkably complex tasks, including associative learning, context-sensitive decision-making, and cross-modal sensory integration (Gibbons et al., 2024; Gibbons, Versace, et al., 2022; Giurfa, 2013; Solvi et al., 2020). Recent studies have also identified brain regions such as the mushroom bodies and central complex that appear to support evaluative processing functionally analogous to that seen in vertebrates (Barron & Klein, 2016; Emanuel & Libersat, 2019; Galili et al., 2014). Yet the question of pain in insects cannot be settled by neural architecture alone. Given the diversity of nervous systems across phyla, and the sheer creative power of adaptive evolution showcased via multiple realisability, behaviour remains our most direct and inclusive route to inferring experience (Elwood, 2011). That is, rather than asking whether an animal has the same hardware, the more relevant question is whether it shows a comparable behavioural profile under similar conditions.

Among insects, much of the extant behavioural evidence concerns bees. Honeybees (*Apis mellifera*) and bumblebees (*Bombus terrestris*) display hallmarks of pain-like states under the Crump et al. (2022) framework: they form associative memories linking noxious stimuli to conditioned cues (Vergoz et al., 2007), regulate foraging based on trade-offs between reward and harm (Gibbons, Versace, et al., 2022), and perform localised self-grooming at injured body parts (Gibbons et al., 2024). These behaviours are persistent, targeted, and sensitive to internal state—features that argue against stereotypy and in favour of evaluative processing. More broadly, there is growing evidence that insects exhibit affective processes beyond pain, ranging from optimism–pessimism biases to play-like behaviour (M. Bateson et al., 2011; Galpayage Dona et al., 2022; Perry & Baciadonna, 2017; Triphan et al., 2025). Such findings suggest that insects can generate and regulate a wider suite of valenced states than once assumed. Still, the bulk of evidence remains taxonomically narrow. In a recent review, Gibbons et al. (2022) assessed six major insect orders and found that while Diptera and Blattodea satisfied multiple pain-related criteria, Orthoptera—grasshoppers, locusts, and crickets—remain underrepresented.

This gap is significant. Orthopterans are both ecologically and evolutionarily important, and central to the edible insect industry: *Acheta domesticus*, the house cricket, is among the most widely farmed insects worldwide, with billions reared annually (Rowe et al., 2024). Orthopterans also represent one of the more basal branches of the winged insects, diverging early within the Polyneoptera (Tihelka et al., 2021). Evidence of sophisticated processing in this group therefore bears directly on questions of how widespread the capacity for felt states may be across insects. Once thought too simple to support sophisticated processing, orthopterans have since been shown to possess nociceptive ion channels (Ylla et al., 2021) and evidence of sensory integration in grasshopper brains (Gupta & Stopfer, 2012). Behavioural data, however, remain scarce. Only two studies have directly tested associative learning with noxious stimuli in this group, both indicating that grasshoppers can learn to avoid harm through limb movement (Forman, 1984; Punzo, 1980), while a third demonstrated crickets’ ability to learn to use visual cues to seek relief from noxious heat (Wessnitzer et al., 2008). Indirectly related efforts have shown behavioural effects of morphine, which increase individuals’ latencies to escape from heated substrates in a dose- and time-dependent manner (Zabala & Gómez, 1991). Given the scale of their use and the ethical stakes of pain, further targeted behavioural tests are urgently needed. Without them, welfare assessments will remain speculative, and our ability to prevent unnecessary suffering limited (Birch et al., 2021; Broom, 2011).

Here we tested a key behavioural criterion for pain in the house cricket *Acheta domesticus*: flexible self-protection, expressed as persistent and directed grooming of a specific site after noxious stimulation (Crump et al., 2022). Self-protection is considered indicative because it implies more than an unconscious withdrawal reflex — the animal must localise the site of injury, sustain attention to it, and direct a motor response accordingly, suggesting some form of body-map awareness. Additionally, the persistence of grooming beyond the initial stimulus is consistent with endogenous modulation of the nociceptive signal, since a purely reflexive response would be expected to terminate with stimulus offset. To that end, we applied a noxiously heated, innocuous, or no stimulus to a single antenna and measured whether crickets groomed the treated antenna more frequently, for longer, and with a distinct temporal profile. If pain-like experience underlies protective grooming, then theory predicts that noxious stimulation should elicit greater, more sustained, and more targeted grooming than either tactile or control conditions.

## Methods

### Study System

We used commercially reared *Acheta domesticus* (house crickets), sourced from Petstock (www.petstock.com.au), and housed them in a laboratory at The University of Sydney, Camperdown, Sydney. We housed crickets in shared containers (40 × 40 × 100 cm) under a 12:12 hour light–dark cycle at 23 ± 1 °C and provided them with free access to wheatgerm and peaches in juice for food and hydration. We conducted all trials under artificial lighting between 10:00 and 15:00 maintaining an ambient light level of approximately 1000 and 100 lux in the higher and lower stress test environments, respectively (detailed below).

We designed this experiment to test whether crickets exhibit flexible, site-specific self-protective behaviour consistent with an evaluative response to noxious stimuli. Our design was broadly inspired by Gibbons et al. (2024), who tested self-directed grooming responses in bumblebees. To examine how environmental context influences such behaviour, we included an additional manipulation comparing responses under higher- and lower-stress conditions; an approach motivated by existing evidence that ambient stressors can modulate the expression of affective behaviour (P. Bateson, 1991).

### Experimental Treatments

We tested 80 adult crickets (40 males, 40 females) in a fully within-subjects design. We assigned each individual to one of two housing conditions, higher-stress or lower-stress (N = 40 each), and exposed them to three treatment conditions in an order randomised via uniform draw (in R): a noxious stimulus (heated probe), a innocuous stimulus (unheated probe), and a control condition involving handling only. The innocuous and control conditions served as procedural and restraint controls, respectively.

For each trial, we gently immobilised the cricket using a sponge and contacted an antenna chosen at random (as above), with side not predetermined or counter-balanced across trials. In the noxious condition, we applied a soldering iron (Hakko FX-888D, 6 mm chisel tip) heated to 65 °C for five seconds. In the innocuous condition we applied the same probe without heating. While in the control condition we handled the cricket in the same manner but did not apply any stimulus. A focal antenna was nevertheless randomly designated for each control trial, providing a chance-level baseline against which grooming allocation in the stimulated conditions could be compared. We selected the temperature of 65 °C in our noxious treatment to provide a stimulus hot enough to be noxious, while avoiding lasting tissue damage. This set point also reflects that used in a comparable test with bumblebees, where it elicited robust self-protective responses without causing visible injury (Gibbons et al. 2024). We confirmed probe temperature prior to each application using a digital infrared thermometer (Sovarcate HS960D). After the application of a given treatment, we immediately transferred the cricket into the observation arena.

We constructed two environmental contexts to coarsely manipulate background stress levels. In the ‘higher-stress’ condition, we placed crickets into open-top plastic arenas (30 × 22 × 10.5 cm) wrapped in red cellophane on the base and sides. This design removed cover, limited substrate for digging, and supplied strong illumination, which elevates vigilance and stress in orthopterans (Hedrick, 2000). In the ‘lower-stress’ condition, we used closed-top arenas with a sand substrate and we reduced ambient light by enclosing them in a brown paper shelter (Fig. 1), thereby providing structural refuge and a more naturalistic, darkened environment. We balanced sex and treatment order across environmental conditions. We filmed all behavioural trials for ten minutes using a Canon 1500D camera fit with a Canon 18-55 mm lens (fixed at 18 mm) at 30 fps, and allowed a ten-minute intertrial interval during which we returned each cricket to its individual holding container.

**Figure 1.**
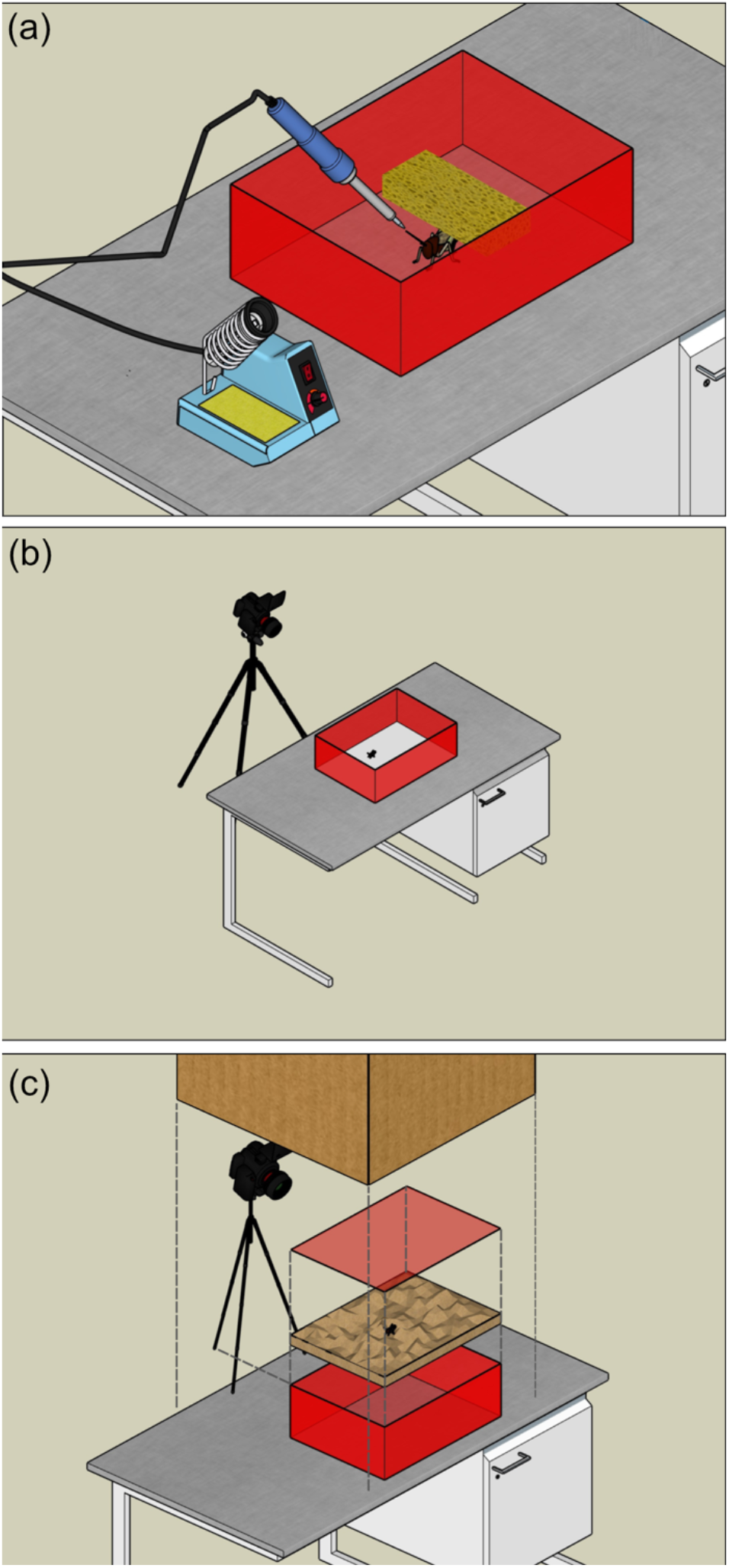
Experimental setup for treatment application and environmental conditions. (a) Noxious treatment: contact of a heated soldering iron tip (65 °C) with a randomly selected antenna; innocuous treatment: contact with an unheated tip; control: no contact applied. (b) ‘Higher-stress’ condition: open-top plastic container (30 × 22 × 10.5 cm) wrapped with red cellophane on the base and sides. (c) ‘Lower-stress’ condition: closed-top, cellophane-wrapped container with a sand layer on the arena floor, placed within a sheltered brown paper structure.

### Behavioural analysis

Three treatment-blind observers coded all grooming behaviour from video recordings using predefined criteria. We defined grooming as either a “touch” (when a cricket brushed the antenna using the prothoracic leg) or “to mouth” (when it drew the antenna toward the mandibles using the same leg), and we recorded which of their two antenna was groomed. Observers recorded timestamps for the start and end of each grooming bout to calculate both frequency and duration. A breakdown of grooming by action type and treatment is provided in Table S1.

### Statistical analysis

We ran four analyses to test our core predictions. First, we asked whether crickets allocated a greater proportion of grooming time to the antenna that had been stimulated. Second, we tested whether they were more likely to groom the stimulated antenna at all. Third, we asked whether grooming of the stimulated antenna lasted longer under noxious conditions than under control or innocuous contact. Fourth, we examined how the temporal profile of grooming evolved across the trial under different treatment conditions.

To test whether crickets preferentially directed grooming to the treated antenna within trials, we fit a beta regression model with the proportion of total antennal grooming duration allocated to the focal side as the response. We included treatment (noxious, innocuous, control), environmental condition (higher-stress vs. lower-stress), and sex as fixed effects, and included individual ID as a random intercept. To satisfy model assumptions, we rescaled proportions to fall strictly within the (0,1) interval. We initially tested a more complex model with a treatment × environment interaction, but this did not improve model fit (χ^2^ = 0.59, df = 2, p = 0.744), so we retained the simpler additive model to maximise power. We then used one-sided post hoc tests on the estimated marginal means to assess whether grooming under each treatment condition was significantly biased toward the stimulated side. We also confirmed that antenna side (left vs. right) had no significant effect in any model (not shown, but see included analysis code), consistent with successful randomisation. We used one-sided tests here, and here only, because the prediction was strictly directional: if noxious stimulation elicits site-directed self-protection, grooming should be biased toward the stimulated antenna, with no theoretical basis for predicting a bias away from it. All other comparisons in this study are two-sided. We report means and confidence intervals on the response scale.

To test the probability of any grooming being directed toward the treated antenna, we used a binomial GLMM. The binary response was coded as 1 if the focal antenna was groomed and 0 otherwise. Fixed effects included treatment, environment, and sex, with individual ID as a random effect. We also tested random slopes for treatment by individual in all models; these either failed to converge or produced near-singular fits, so we retained the random-intercept specification (D. J. Barr et al., 2013). A full model including the treatment × environment interaction again showed no improvement in fit (χ^2^ = 1.22, df = 2, p = 0.544) and was simplified accordingly.

To test the duration of directed grooming, we used a Tweedie GLMM with a log link, which allowed us to model the right-skewed data with many zero values. The model included treatment, environmental condition, and sex as fixed effects, with individual ID as a random intercept. The more complex model including the treatment × environment interaction did not improve fit (χ^2^ = 0.37, df = 2, p = 0.830) and was likewise simplified.

Finally, to explore the temporal dynamics of grooming, we fitted generalised additive models (GAMs) to the durations of focal grooming bouts over the ten-minute trial. Each observation in the temporal model corresponded to a single focal grooming bout, with bout onset time (seconds into the trial, from video timestamps) as the continuous predictor and bout duration as the response. Each model included treatment as a parametric term, treatment-specific smooths for time since stimulus, and a random intercept for individual identity. We used a Tweedie distribution with a log link to accommodate zero-inflated, right-skewed bout durations. Predictions were generated marginal over individuals by excluding the random-effect term, and we back-transformed estimates to the response scale with 95% confidence intervals derived from the standard errors. These models allowed us to visualise whether grooming in response to noxious stimulation was transient or sustained across time, and whether its temporal profile differed significantly from control and tactile treatments.

We conducted all statistical analyses in RStudio (v4.3.2; RStudio Team, 2024), using the packages lme4 for mixed models (Scheipl, 2011), glmmTMB for Tweedie models (Brooks et al., 2017), mgcv for GAMs (Wood, 2018), DHARMa for residual diagnostics (Hartig, 2022), and emmeans for post hoc comparisons (Russell et al., 2018). We adopted the conventional threshold of statistical significance at α = 0.05, and report all summary statistics with their standard errors.

### Ethical note

Although invertebrate research is not subject to formal ethics approval in Australia, we followed high standards of care throughout. We kept crickets in stable environmental conditions, provided continuous access to food and water, and ensured that no procedures caused injury or lasting harm. After completing the study, we returned all individuals to their housing containers, where they lived out their natural lifespans under continued laboratory care with free access to food and water.

### Data and code availability

All underlying data and code are freely available via GitHub (https://github.com/invertBEACON/cricket_protection) and will be persistently archived upon acceptance.

## Results

### Specificity of grooming

Within trials, crickets allocated a significantly greater proportion of antennal grooming to the stimulated side following noxious treatment compared to control (β = 0.62 ± 0.29, *z* = 2.14, *p* = 0.032; Fig. 2a; Table 1). On average, 59% (95% CI: 0.505–0.674) of grooming time was directed to the focal antenna after noxious stimulation, compared with 44% (0.337–0.548) in no-contact control trials and 52% (0.418–0.615) in innocuous contact trials. A one-sided test against chance confirmed that the noxious proportion was significantly greater than 0.5 (*z* = 2.07, *p* = 0.019), while no-contact control and innocuous treatments did not differ from chance. There was no effect of environmental condition (β = −0.14 ± 0.23, *z* = −0.62, *p* = 0.538) or sex (β = −0.37 ± 0.24, *z* = −1.56, *p* = 0.118).

**Figure 2.**
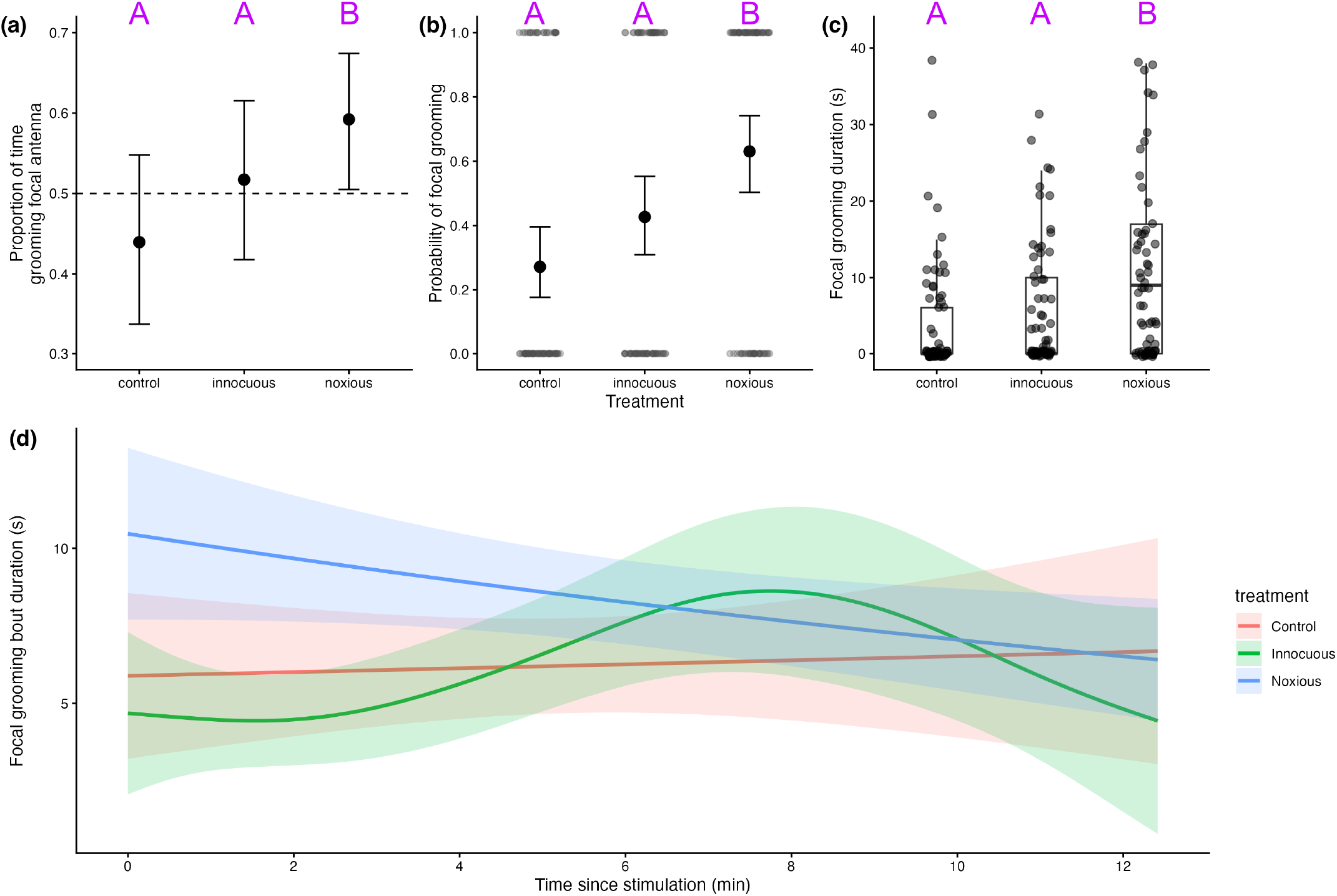
Directed grooming responses following antennal stimulation in crickets. (a) Proportion of grooming time allocated to the stimulated antenna, shown as estimated marginal means (± 95% CI) from a beta regression. (b) Probability that the stimulated antenna was groomed at all, with grey points indicating individual responses (1 = groomed, 0 = not groomed) and black points ± 95% CI from a binomial GLMM. (c) Duration of grooming directed to the stimulated antenna, with dots representing individuals and boxplots showing distributions by treatment. (d) Temporal dynamics of focal-antennal grooming over the 12-min trial, with lines depicting treatment-specific smooths from a GAM and shaded ribbons showing standard errors. Letters above plots (a) through (c) denote grouping by statistical significance of post-hoc tests.

**Table 1:** Summary of model results for directed grooming responses following antennal stimulation. Results are presented for three models: (i) the proportion of grooming directed to the stimulated antenna (beta regression), (ii) the probability that any grooming was directed toward the stimulated antenna (binomial GLMM), and (iii) the duration of directed grooming bouts (Tweedie GLMM). Fixed effects include treatment (control, innocuous contact, noxious heat) and environmental condition (higher-vs. lower-stress); all models included individual ID as a random effect. Estimates (est.) are shown with standard errors (s.e.), test statistics (z), and *p*-values. Treatment contrasts are relative to the control condition, which serves as the reference level (intercept).

| Effect | Est. | SE | z | p |
| --- | --- | --- | --- | --- |
| *Specificity of grooming* |  |  |  |  |
| intercept | -0.24 | 0.26 | -0.94 | 0.35 |
| treatment (innocuous) | 0.33 | 0.30 | 1.10 | 0.27 |
| treatment (noxious) | 0.68 | 0.28 | 2.40 | **0.016** |
| environment (higher-stress) | -0.12 | 0.23 | -0.54 | 0.59 |
| sex (male) | 0.01 | 0.23 | 0.04 | 0.97 |
| *Grooming probability* |  |  |  |  |
| intercept | -1.53 | 0.40 | -3.8 | **< 0.01** |
| treatment (innocuous) | 0.69 | 0.36 | 1.91 | 0.06 |
| treatment (noxious) | 1.52 | 0.38 | 3.96 | **< 0.01** |
| environment (higher-stress) | 0.66 | 0.34 | 1.95 | 0.051 |
| sex (male) | 0.42 | 0.34 | 1.25 | 0.21 |
| *Grooming duration* |  |  |  |  |
| intercept | 1.10 | 0.31 | 3.52 | **< 0.01** |
| treatment (innocuous) | 0.60 | 0.30 | 2.03 | **0.04** |
| treatment (noxious) | 1.39 | 0.27 | 5.14 | **< 0.01** |
| environment (higher-stress) | 0.10 | 0.24 | 0.43 | 0.67 |
| sex (male) | 0.11 | 0.24 | 0.46 | 0.64 |

### Directed-grooming probability

Crickets were significantly more likely to groom the antenna that received the noxious stimulus compared to the no-contact control (log-odds = 1.52 ± 0.38, *z* = 3.96, *p* < 0.001), with grooming also elevated—but not significantly so—under the innocuous treatment (log-odds = 0.69 ± 0.36, *z* = 1.91, *p* = 0.056). On the response scale, we estimated that grooming occurred on the focal side in 63% of noxious trials (95% CI: 0.50–0.74), 43% of innocuous trials (0.31–0.55), and 27% of control trials (0.18–0.40). Post hoc comparisons confirmed a robust noxious–vs-no-contact control difference (Δlog-odds = −1.52 ± 0.38, *z* = −3.96, *p* < 0.001), with weaker support for a noxious–vs-innocuous difference (Δlog-odds = −0.83 ± 0.36, *z* = −2.34, *p* = 0.051) and no significant difference between innocuous and no-contact control (Δlog-odds = −0.69 ± 0.36, *z* = −1.91, *p* = 0.136). While there was a trend toward increased grooming under the higher-stress condition (β = 0.66 ± 0.34, *p* = 0.051), this was not statistically significant,

### Directed-grooming duration

Crickets groomed for significantly longer following noxious stimulation than under no-contact control (*est*. = 1.39 ± 0.27, *z* = 5.15, *p* < 0.001) or innocuous (est.= 0.60 ± 0.30, *z* = 2.04, *p* = 0.042) treatment conditions (Fig. 2c; Table 1). The average estimated grooming duration was 13.48 ± 2.39 s after noxious treatment, compared to 6.12 ± 1.32 s after innocuous and 3.35 ± 0.82 s in the no-contact control. Post hoc pairwise comparisons confirmed a significant increase in grooming duration under noxious treatment compared to no-contact control (ratio = 0.25 ± 0.07, *z* = –5.15, *p* < 0.001) and innocuous (ratio = 0.45 ± 0.11, *z* = –3.29, *p* = 0.0029). Grooming duration did not differ significantly between innocuous and no-contact control treatments (ratio = 0.55 ± 0.16, *z* = –2.04, *p* = 0.104). We detected no effect of environmental condition (*est*. = 0.10 ± 0.24, *z* = 0.43, *p* = 0.67) or sex (est. = 0.11 ± 0.24, *z* = 0.46, *p* = 0.64) on grooming duration.

### Temporal dynamics of directed-grooming

We found that including treatment-specific smooth terms significantly improved model fit (*F* = 2.64, *df* = 9.8, residual *df* = 405, *p* = 0.036), confirming that the duration of antennal grooming varied meaningfully over time in a treatment-specific manner. In plain terms, we detected clear treatment effects on the timecourse of directed grooming: the noxious stimulus led to sustained, elevated grooming that declined across the observation period; innocuous stimulation produced a delayed peak in grooming around minute 8; and no-contact control responses remained stable and low throughout (Fig. 2d).

Smooth terms were statistically significant for both noxious (*edf* = 1.00, *F* = 6.996, *p* = 0.009) and innocuous (*edf* = 3.39, *F* = 2.813, *p* = 0.024) treatments, indicating reliable temporal variation in grooming under those conditions. In contrast, grooming under no-contact control conditions showed no significant change over time (*edf* = 1.00, *F* = 0.003, *p* = 0.954), consistent with a flat response.

Together, these results support our prediction that noxious stimulation elicits both more frequent and more persistent grooming — a behavioural profile suggestive of flexible, experience-guided self-protection. The innocuous treatment also produced an extended grooming response, though with a delayed and less consistent timecourse, potentially indicating residual mechanical irritation or procedural artefacts.

## Discussion

Pain remains one of the most elusive and consequential frontiers in animal cognition, and insects provide a particularly demanding test case. As outlined above, behavioural evidence— particularly flexible, site-specific responses to harm—offers our most direct route to inferring pain-like states in these animals (Crump et al., 2022). Our findings demonstrate such a response in the evolutionarily and commercially significant *Acheta domesticus*: crickets groomed a noxiously stimulated antenna more frequently, for longer, and with a distinct temporal profile compared to tactile or no-contact controls. The response was both site-specific and persistent, suggesting that crickets monitor injury location and adjust their behaviour in ways not reducible to simple reflexes.

Crickets’ heightened grooming intensity followed a clear temporal trajectory (Fig. 2c): noxiously stimulated individuals showed an elevated, sustained phase that gradually declined, a pattern reminiscent of findings in bees and rodents (Gibbons et al., 2024; Mittal et al., 2009). By contrast, the innocuous contact treatment produced an early dip in grooming before a later rise. One possibility is that sudden, non-noxious touch was initially treated as a potential predatory threat, eliciting vigilance or an orienting response that temporarily displaced grooming (Zacarias et al., 2018). Alternatively, the tactile stimulus may have prompted competing behaviours such as scanning or locomotion, which delayed antennal maintenance until the salience of the stimulus subsided (Oram & Card, 2022). Similar context-dependent processing has been demonstrated in crickets: when antennae were contacted with a glass rod, a hunting spider leg, or an orb-weaver leg, responses varied from aggression to aversion to search behaviour, with glass-rod contact especially likely to elicit search or no response (Okada & Akamine, 2012). This parallel suggests that our probe, like the glass rod, was interpreted flexibly rather than as a simple mechanical trigger. Overall, the observed patterns align with the criterion of flexible self-protection and add Orthoptera to the short list of insect orders for which such evidence exists (Gibbons, Crump, et al., 2022).

We also manipulated environmental stress to test its influence on behavioural expression, given prior evidence that stress can suppress affectively mediated responses (Perry & Baciadonna, 2017). Grooming likelihood and duration did not differ significantly between ‘higher-’ and ‘lower-stress’ contexts, and we likewise found no effect of sex in any model. This may indicate that our environmental manipulation lacked sufficient salience, and/or that self-protective responses are robust to background stress and consistent across sexes. Either way, the persistent expression of grooming across environmental and sex conditions supports the view that these behaviours are internally regulated and functionally significant, rather than fragile or reflexively triggered.

Taken together, our findings further erode the once-dominant view of insects as behaviourally rigid and insentient. Crickets have already been shown to possess nociceptors (Ylla et al., 2021), centralised sensory integration (Gupta & Stopfer, 2012), and have the capacity to learn from aversive events (Forman, 1984; Punzo, 1980), and moderate their responses to noxious stimuli following analgesia (Zabala & Gómez, 1991). By demonstrating flexible self-protection, we add a fifth pillar to this growing body of evidence in Orthoptera and strengthen coverage of the core criteria for pain-like states outlined by Crump et al. (2022). Together, these lines of evidence suggest that orthopterans exhibit the same conjunction of features—nociception, integrative processing, learning, and targeted self-protection—that are taken to warrant serious consideration of sentience. Similar convergence in decapod crustaceans, including site-directed grooming in glass prawns modulated by local anaesthesia (S. Barr et al., 2008; S. Barr & Elwood, 2024), and aversive responses to chemical irritants in shore crabs (Elwood et al., 2017), has already contributed to their legal recognition as sentient in some jurisdictions (e.g. the UK Animal Welfare (Sentience) Act 2022). While the current evidence base does not collectively resolve the question of insect pain, it meaningfully narrows the evidential gap and underscores the need for further comparative and taxonomically broad investigations.

The implications of this work extend beyond theory. *Acheta domesticus* is reared by the millions for food, feed, and research, often under conditions that assume an absence of felt experience. If insects can respond to injury in a way consistent with pain—as our findings suggest—then the ethical landscape shifts. Under a precautionary approach (Birch, 2024), the burden moves toward reducing harm in farming, handling, and experimentation. This shift is not only moral, it reflects the accelerating recognition, across fields, that sentience may be evolutionarily deep and taxonomically widespread.

The expanding frontier of behavioural evidence is not just pushing disciplinary boundaries, it is redrawing ethical ones. As more insect taxa exhibit traits once thought exclusive to vertebrates, the phylogenetic perimeter of likely sentience continues to creep. Our findings join a broader movement toward reassessing long-held assumptions, urging not premature conclusions but a recalibration of care in the face of uncertainty; one that may prove transformative for how we regard the smallest of animals.

## Acknowledgements

This work was supported by the Australia and Pacific Science Foundation (APSF240014). We thank Professor Elwood and one anonymous reviewer for their thoughtful feedback and suggestions.

