## Supplementary material for "Flexible self-protection as evidence of pain-like states in house crickets"

**Table S1:** Breakdown of focal grooming by action type (leg touch vs. to-mouth) and treatment. Values are computed from grooming bouts directed to the stimulated (focal) antenna only. To-mouth grooming dominated across all treatment conditions, with the relative proportion of action types remaining broadly consistent.

| **Treatment** | **Action** | **N bouts** | **Total dur. (s)** | **Mean dur. (s)** | **SD (s)** |
| --- | --- | --- | --- | --- | --- |
| Control | To mouth | 24 | 208 | 8.67 | 4.48 |
| Control | Touch | 16 | 61 | 3.81 | 4.18 |
| Innocuous | To mouth | 53 | 467 | 8.81 | 5.21 |
| Innocuous | Touch | 18 | 46 | 2.56 | 3.07 |
| Noxious | To mouth | 86 | 879 | 10.2 | 5.03 |
| Noxious | Touch | 39 | 217 | 5.56 | 6.46 |
